# A python-based package for long-lasting video acquisition and semi-automated detection of convulsive seizures in rodents

**DOI:** 10.1101/2022.04.15.488472

**Authors:** Daniel Diaz-Arce, Anis Ghouma, Paolo Scalmani, Massimo Mantegazza, Fabrice Duprat

**Affiliations:** Université Côte d’Azur, Valbonne-Sophia Antipolis, France; U.O. VII Clinical Epileptology and Experimental Neurophysiology, Foundation IRCCS Neurological Institute Carlo Besta, Milan, Italy; CNRS UMR7275, Institute of Molecular and Cellular Pharmacology (IPMC), Valbonne-Sophia Antipolis, France; Inserm, Valbonne-Sophia Antipolis, France

**Keywords:** Epilepsy, image analysis, noninvasive, cost-effective, movement tracking, convulsion

## Abstract

**Background:** Epilepsy is a very invalidating pathology characterized by the unpredictable appearance of abnormal cerebral activity leading to seizures and co-morbidities. The ability to detect and even predict seizures is a major challenge and many research laboratories are using rodents’ models of epilepsy to unravel possible mechanisms. The gold standard to record and detect seizures is electroencephalography, but it is very invasive. For rodents used in research, video analysis is a very interesting approach but the major disadvantages are that it is time consuming, prone to human error, and not very reproducible. Commercial solutions for detailed phenotyping analysis on humans or rodents exist but they are costly. Some open source software programs are also available, they provide very interesting and precise behavior data, but none of them are made for high throughput analysis of a large number of video files generated by long lasting recordings.

**New method:** We developed an open-source python-based package of two software programs that enable automated video acquisition and simple motion analysis associated with a spectral power analysis, which enable a semi-automated identification of convulsive seizures. The method needs cheap webcams and a computer or a server.

**Results:** Using two murine epilepsy models (Nav1.1 mutations), we have compared our motion analysis software to human visual inspection and found an 88.8% accuracy in convulsive seizures detection. We then compare our method to the gold standard electrocorticogram analysis and found a 93.2% accuracy. The motion analysis is also interesting to get a readout of the animal activity without the invasiveness of electromyogram recordings.

**Conclusions:** This new method is easy to use, cost-effective and allows: 1) detection of convulsive seizures in a noninvasive way, 2) high speed analysis of a large number of video files with a good accuracy, and 3) automated acquisition and semi-automated analysis of a very large number of files.

**Highlights:**

- Noninvasive semi-automated detection of convulsive seizures from videos
- High speed of analysis with a good accuracy
- Ability to acquire and analyze a very large number of files
- Easy to use graphical interfaces
- Cost-effective setup

## 1 Introduction

Epilepsy is a very invalidating pathology and the fourth most common neurological disease (Fisher, Acevedo et al. 2014). It is an episodic disorder characterized by the unpredictable appearance of abnormal cerebral activity leading to recurrent seizures and by a wide range of co-morbidities (Scheffer, Berkovic et al. 2017). Although many anti-epileptic drugs have been commercialized in the last decades, about one-third of people with epilepsy are still pharmacoresistant (Loscher and Schmidt 2011). Many research laboratories study the mechanisms leading to the pathological state (chronic epilepsy), called epileptogenesis, and the mechanisms triggering seizures, called ictogenesis (Pitkänen 2017). The use of rodent models of epilepsy enables to shed light on these two mechanisms and to test new anti-epileptic treatments; the evaluation of these treatments on spontaneous seizures of models showing chronic epilepsy can greatly improve the translational value of the study (Loscher and Schmidt 2011). Epileptic seizures occur at variable frequencies, implying long term recordings (over days or weeks) for an accurate quantification, which is essential for the characterization of models and the assessment of the effects of treatments on spontaneous seizures. The features of seizures have been precisely classified both in humans (Fisher, Cross et al. 2017) and in rodents (Racine 1972, Velíšková 2017), but the ability to detect seizures with automated or semi-automated methods is still a major challenge (Giannakakis 2014).

The gold standard for recording and detecting seizures is electroencephalography, but it needs neurosurgery for the implantation of invasive electrodes or the use of a helmet with scalp electrodes, which are uncomfortable for humans and not possible to be used on mice for long term recording. There is a growing interest in movements and motion analysis for detecting seizures and behavioral features in general. For humans, the use of contact sensors (accelerometers, magnetic sensors, electromyography, piezoelectric sensors) enables a detailed analysis with a moderate discomfort, but their use is very difficult on rodents because of the small size of the animals. The use of sensors without contact, like video cameras and radars, is thus a growing field of research and commercial development. For humans, many commercial systems that use these sensors are proposed, and they are often associated with contact sensors. The sensitivity of detection of tonic-clonic seizures of these systems varies from 31% to 95% (Arends 2018). For rodents used in research, video analysis is a very interesting approach, because it is not invasive and enables to record animals in their home cage at any age. Many studies have been performed by manual annotation of long duration videos, but the major disadvantages are that this approach is 1) very time consuming, 2) prone to human error, and 3) not reproducible because two different annotators usually do not agree at 100%.

Commercial solutions for detailed phenotyping analysis on humans or rodents exist, e.g. Noldus (http://www.noldus.com, accessed the 1/6/2021), CleverSys (http://cleversysinc.com, accessed the 1/6/2021), MouseSpecifics (http://mousespecifics.com, accessed the 1/6/2021), or ANY-maze (http://www.anymaze.co.uk, accessed the 1/6/2021). They provide a detailed phenotypic analysis but they are expensive and most of the videos and analysis files are produced in a proprietary format that needs extra export manipulations to be used with other programs (when they are available).

Some open source software programs are also available. MouseMove is based on the commercially available LabVIEW development system and gives information like travel distance, speed, turning and curvature for mice during an open field behavioral test (Samson, Ju et al. 2015). Another project based on commercially available Matlab language enables very precise behavioral phenotyping (e.g. drinking, eating, grooming, …) using a video database as reference (Jhuang, Garrote et al. 2010). Some limitations are that the system needs high contrast (dark mice on a white background), recording from the side view, and process one cage at a time. Another project for studying neurodegenerative disorders presents a high precision motion analysis using 6 cameras, markers fixed on the animal, and a constrained walkway (Karakostas, Hsiang et al. 2014). The Live Mouse Tracker project allows to track and precisely monitor the activities of 4 mice using implanted radio-frequency identification (RFID) methods (de Chaumont, Ey et al. 2019). The most efficient available software programs use deep learning methods and enable marker-less motion capture (reviewed in Mathis and Mathis 2020). These software programs give invaluable pose estimation measurements but need computational power (most of them use Graphical Processing Unit, GPU). For example, DeepLabCut can process low resolution video (204×162 pixels) at around 85 Hz (Nath, Mathis et al. 2019).

All these systems can provide very interesting and precise behavioral data, but none of them is designed for high throughput analysis of a large number of video files generated by long lasting recordings. Moreover, the available systems have not been developed and thoroughly tested for identifying seizures. Thus, there is a need for a simple software to acquire videos from a large range of cheap IP cameras, perform a quick and intuitive analysis on a standard computer, in particular for identifying seizures, and produce data in common file formats. This is why we developed a package of two software programs that enable video acquisition and motion analysis, VASD (Video Acquisition and Seizure Detection) and SASDI (Semi-Automatic Seizure Detection on Images), which we used to rapidly identify convulsive seizures, which are characterized by a sudden rise in motion amplitude and frequency power of the videos.

## 2 Materials and methods

### 2.1 Video-ECoG-EMG acquisition

Mice were implanted with ECoG electrodes (mouse stainless steel screws, PlasticsOne, USA) in anterior locations on each hemisphere and with EMG electrodes (stainless steel wire, Plastics Ones) at P32 under isoflurane anesthesia. EMG wires were inserted in the trapezius (neck) muscles. The reference electrode was implanted in the occipital bone above the cerebellum. To stabilize the connectors, 2 screws were fixed to the skull with a resin (Super Bond, Sun Medical, Japan); wires and the connector (PlasticsOne, USA) were fixed with dental cement (Dentalon, Kulzer, Germany). Mice received anti-inflammatory drug for 3 days (Carprofen 0.04 mg/mL in drinking water, KRKA, Slovenia) for a better recovery. After recovering from surgery, animals were allowed to rest for 3 days. They were placed in individual recording cages (half a cage for each animal) and connected to an amplifier (Animal Bio Amp, AD Instruments) through a cable and a swivel to allow the mouse to freely move in the cage. The amplifiers were connected to an acquisition system (PowerLab 16/35 and Labchart 8, AD Instruments). A video of each pair of animals was acquired with a batch of 4 analogic cameras (ZC-YX270PE, Ganz) connected to a computer (Precision, Dell) through a video multiplexer (VM-HD4, cctvcamerapros) and a video digitizer (Dazzle video recorder HD, Pinnacle), allowing the simultaneous recording of 8 animals.

### 2.2 Acquisition of videos

We used different brands and types of cameras. The video-EMG cameras described above, a batch of 4 cameras (HDCVI, 2 Mpx, Sony exmor) equipped with infrared light and connected to a digital video recorder (DVR, HDVCI 1080P) having its own IP address within our institute internal network (rate 1 Gbps). A camera with an internal microphone (POE HX, sv3c Shenzhen, China) having also its own IP address.

### 2.3 Computers

For running VASD, we have used a PowerEdge R940 server (Dell, Ireland) with 4 processors (Intel Xeon Gold 5220, total of 72 cores), 64 Go of RAM, and 36 To of hard drive, but it can run on any linux-based computer.

SASDI can run on various laptops (we tested i5 to i7 processors based computers).

### 2.4 Experimental animals

All experiments were performed according to policies on the care and use of laboratory animals of European Communities Council Directive (2010/63EU) and under the agreement number 04551.02 from the French Ministry of research. R1648H knock-in (Scn1a^RH/+^) mice (Martin, Dutt et al. 2010, Hedrich, Liautard et al. 2014, Mantegazza and Broccoli 2019, Salgueiro-Pereira, Duprat et al. 2019, Mantegazza, Cestele et al. 2021) were in a 50/50 % mix genetic background C57Bl/6J and 129/SV; they were induced at P21 with a protocol of chronic hyperthermia (10 days, 1 per day), which leads to chronic epilepsy and co-morbidities mimicking Dravet syndrome (Salgueiro-Pereira, Duprat et al. 2019), and were recorded (synchronized video-ECoG-EMG) between P34 and P60. Heterozygous knock-out *Scn1a* (Scn1a^+/-^) mice are an established model of Dravet syndrome (Yu, Mantegazza et al. 2006, Liautard, Scalmani et al. 2013, Mantegazza and Broccoli 2019, Mantegazza, Cestele et al. 2021) they were in the C57Bl/6J background and were recorded (video) between P21 and P40.

### 2.5 Algorithms

The same module is used by VASD and SASDI to analyze video files, it is based on opencv methods:

1. each frame of the video file is converted to LAB color (L=Lightness, A=green/red, B=blue/yellow) and Lightness channel is filtered with a Contrast Limited Adaptive Histogram Equalization (clip limit 3, grid size (8, 8)) to correct noise and bad shading;
2. the difference of each pixel of consecutive frames is calculated, filtered with a median blurring (aperture size 3) to remove isolated pixels’ noise and thresholded (binary above 25) to remove background noise;
3. the resulting frame is used to calculate the sum of all pixels’ values within each ROI and normalized to the maximal value (number of pixels times 255).

Users should try to use similar ROIs for each animal because the ratio between the size of the animal and the ROI surface influences the calculated motion. This will facilitate the choice of a motion value threshold over which the probability to observe a seizure is high. This threshold will depend mainly on the animal motion variations between normal behavior and during convulsive seizures, but the quality of the video (resolution, bitrate, frame rate, …), its distance from the animal, the intensity of lightning, and the size of the ROI can influence the threshold.

The motion values can also be displayed as a power spectrum index. Using Scipy signal module, a spectrogram is calculated per segments of 1 s without overlap, using a Tukey window (alpha=0.25, symmetrical), then for each segment all frequencies power spectra are summed. The displayed index provide a measurement of cyclic phenomena present in motion, thus convulsive seizure gives a high index in comparison to basal activity, as illustrated in Figure 1A (convulsive seizure is indicated by a red arrow). We also tested some low quality video recordings during night that gives a “snowy” video and produces a large constant artefact in motion analysis. Of note, the seizures are easily identified upon spectral analysis (see Figure 2), showing its usefulness.

**Fig. 1.**
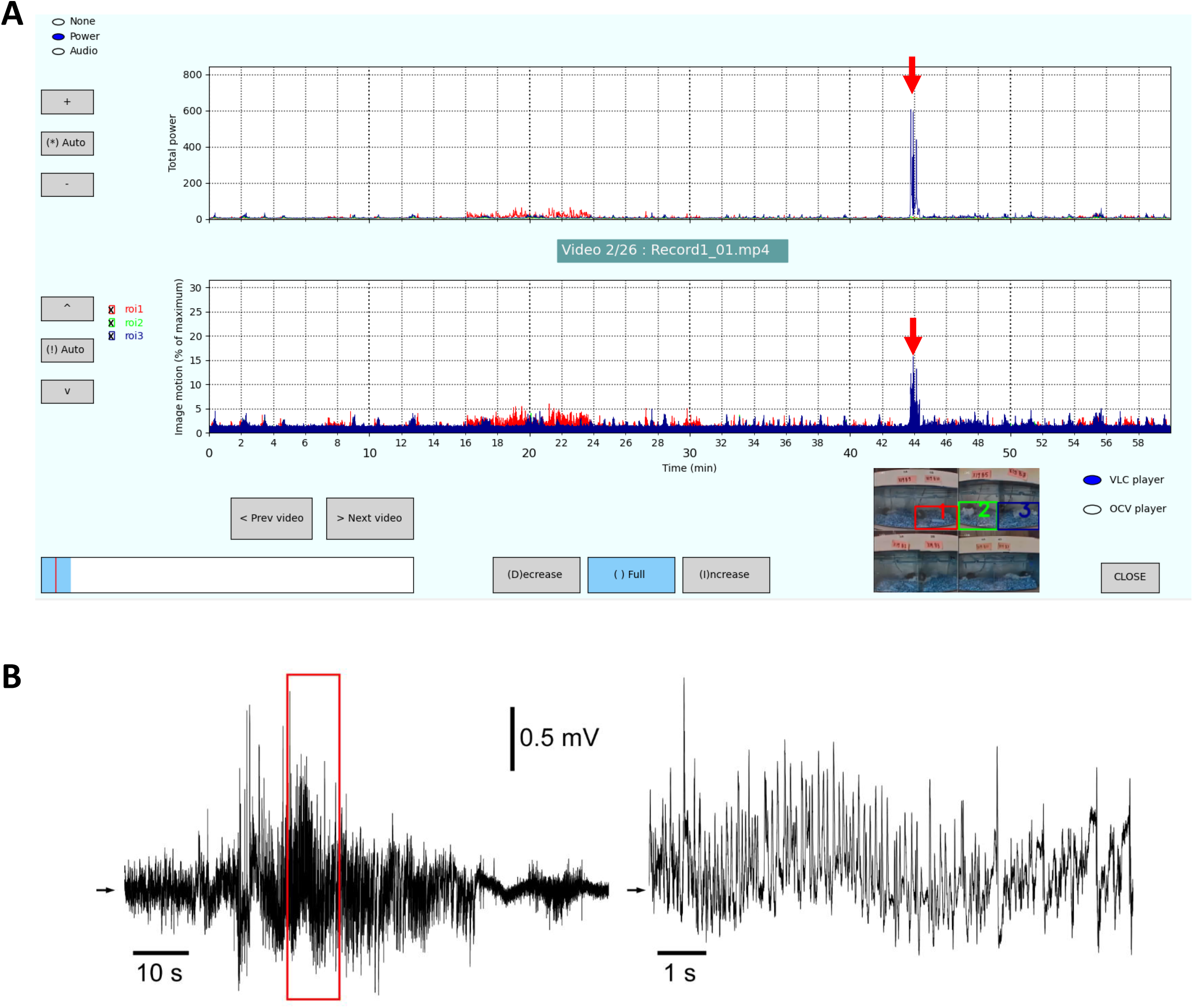
Example of SASDI interface with results of motion analysis. A: Example of the analysis output: “Results” page of SASDI interface. The lower graph is the motion analysis (normalized to maximal motion), here performed on 3 regions of interest (ROI) on the current video (color-coded ROI bounds are displayed on top of video image in the lower right panel). Upper part is displayed optionally, in this example the power spectral index of the motion analysis (sum of all spectral powers) is shown, it can also be the audio stream of the video file, or an empty graph. Various buttons allow the selection of video files and scaling. Clicking on the motion graph opens the analyzed video file at the corresponding time to allow the visual check of the putative seizure. In this example, high motion and power spectral index (red arrows) occurred at around 44 min in the ROI3 (corresponding to the animal in upper right cage on the video). The visualization of the video allowed to confirm that is was a seizure and to classify its severity according to our scale. B: Corresponding ECoG traces showing cortical electrical activity during a convulsive seizure. The parts highlighted by the boxes (left panel) are displayed on the right panel at a higher time resolution. The black arrows indicate 0 mV.

**Fig. 2.**
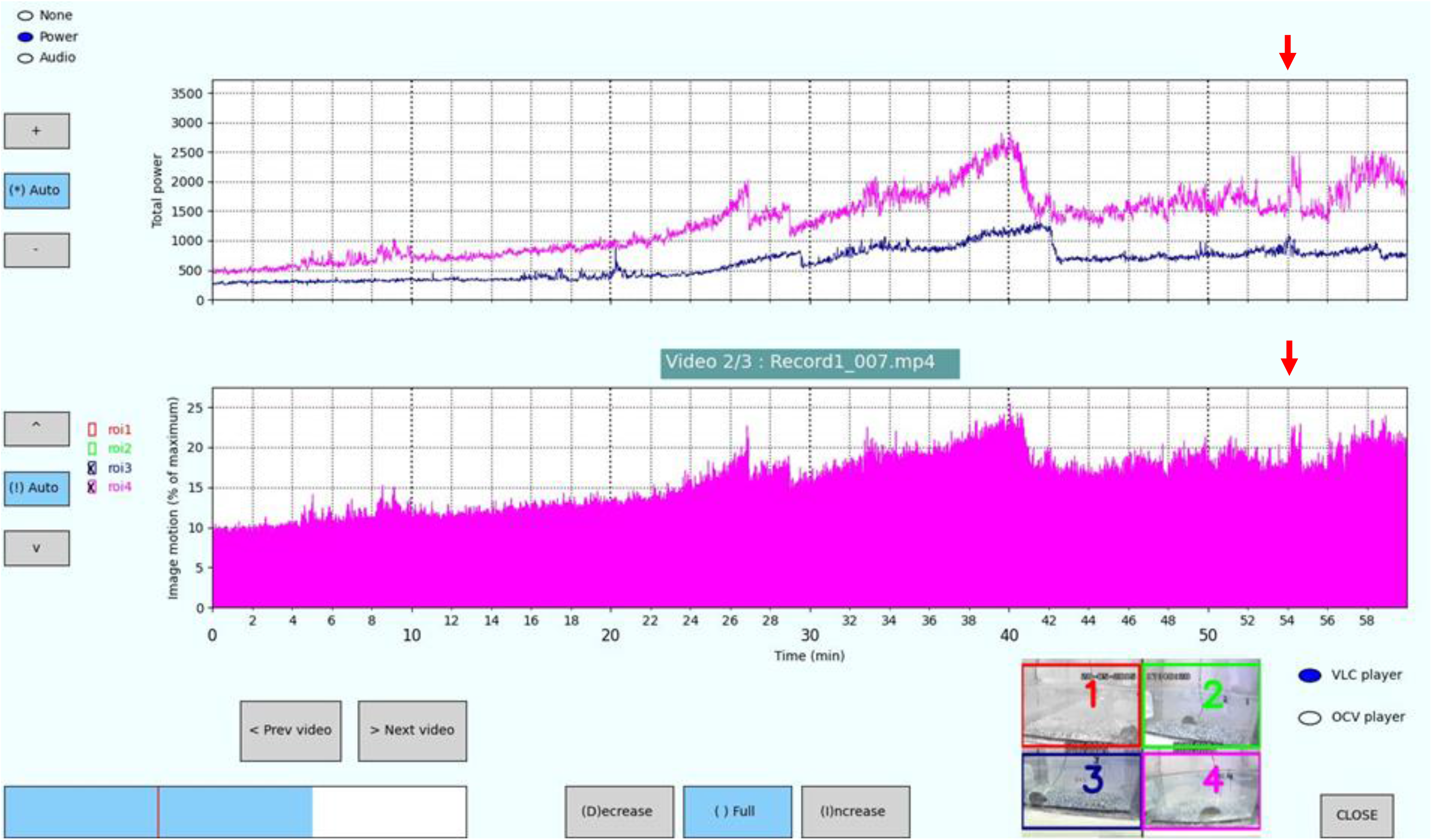
Spectral features enables the analysis of poor quality video. Example of the analysis of a low quality “snowy” video with SASDI. A seizure occurring at around 54min (red arrows) in ROI4 (magenta) can still be spotted mostly with spectral analysis, showing the interest of this approach to help the user to spot seizures (the display of ROI1 and ROI2 are off, ROI3 is hidden by ROI4 in bottom graph).

## 3 Results

### 3.1 Development of a Python based package

Our goal is to avoid the need of costly specialized setups and invasiveness, thus we have chosen to use simple and cost-effective IP cameras or webcams and standard or slightly modified cages.

We chose python language because it can run under an open source license, making it freely usable and distributable (https://www.python.org/psf/, accessed the 1/6/2021). It is largely used in the scientific community, and it is cross-platform (Linux, Windows, Mac, etc.). The video acquisition exploit FFmpeg (developed under GNU General Public License, https://ffmpeg.org/, accessed the 1/6/2021), the leading multimedia framework, which is cross-platform and deals with most of the available codecs, enabling to capture and save most video formats. The image analysis is based on another widely used open source package, opencv (https://opencv.org/, accessed the 1/6/2021), enabling most image transformations and analyses. Finally, these scripts used packages from Python 3.9 standard libraries and from open source python packages like Schedule (https://pypi.org/project/schedule/, accessed the 1/6/2021), Numpy (https://numpy.org), Matplotlib (https://matplotlib.org, accessed the 1/6/2021), and SciPy (https://scipy.org, accessed the 1/6/2021).

Different approaches are possible for motion tracking (Abbas and Masip Rodo 2019). We chose temporal differencing, which computes the pixel-by-pixel difference in consecutive video frames. This implies a non-stationary reference image, which is an important feature for analyses of long term recordings, because the animal surrounding changes with time (bedding, food, etc.) and the light switches from white to infrared during day/night cycles. We also tested approaches based on quantification of optical flow, the pattern of apparent motion obtained comparing successive frames in the video sequence (Abbas and Masip Rodo 2019), but the computational cost of this method is high and we focused on an approach for semi-automated seizure detection on images that can be implemented on most computers.

Our package is divided into two parts:

#### 3.1.1 Automated video acquisition and motion detection

VASD (Video Acquisition and Seizure Detection) is designed to run on a linux server and enables to acquire videos from up to 8 IP cameras using a rtsp stream (exchange format that most IP cameras can provide). VASD acquires series of 1 hour-long video files stored in the mp4 multimedia container format generated using the highly efficient H265 codec. The acquisition duration can be very long, up to months (the only limitation is the hard drive storing space). The acquired video files correspond to camera full field or half fields (left or right parts of the field in separated video files). VASD performs automatically a motion analysis on each file, storing results in a csv format file. VASD analyzes the full acquired video field or both sides separately. VASD uses multiprocessing module that enables the analysis of many videos in parallel, but only one thread is used for each video file because our test demonstrated that using more than one thread decreases the performance. For example, on a Dell server, CPU Intel Xeon Gold 5220, one video (15 frames per seconds, 640×480 pixels) is analyzed at a rate of 61 images/s (Hz) using 4 threads for the same video file, 119 Hz using 2 threads, and 123 Hz using 1 thread. At present, VASD does not work on Windows platforms because the used schedule package is not working properly (does not start at the proper time), but future versions of the package will probably allow to use it on Windows based computers. The motion analysis implemented in VASD is a normalized sum of variations in intensities pixel by pixel (as described in the above Algorithms paragraph). This is performed for each Region of Interest (ROI). Results (motion index and time from video start) are saved to standard csv files. The VASD user interface is shown in Figure 3.

**Fig. 3.**
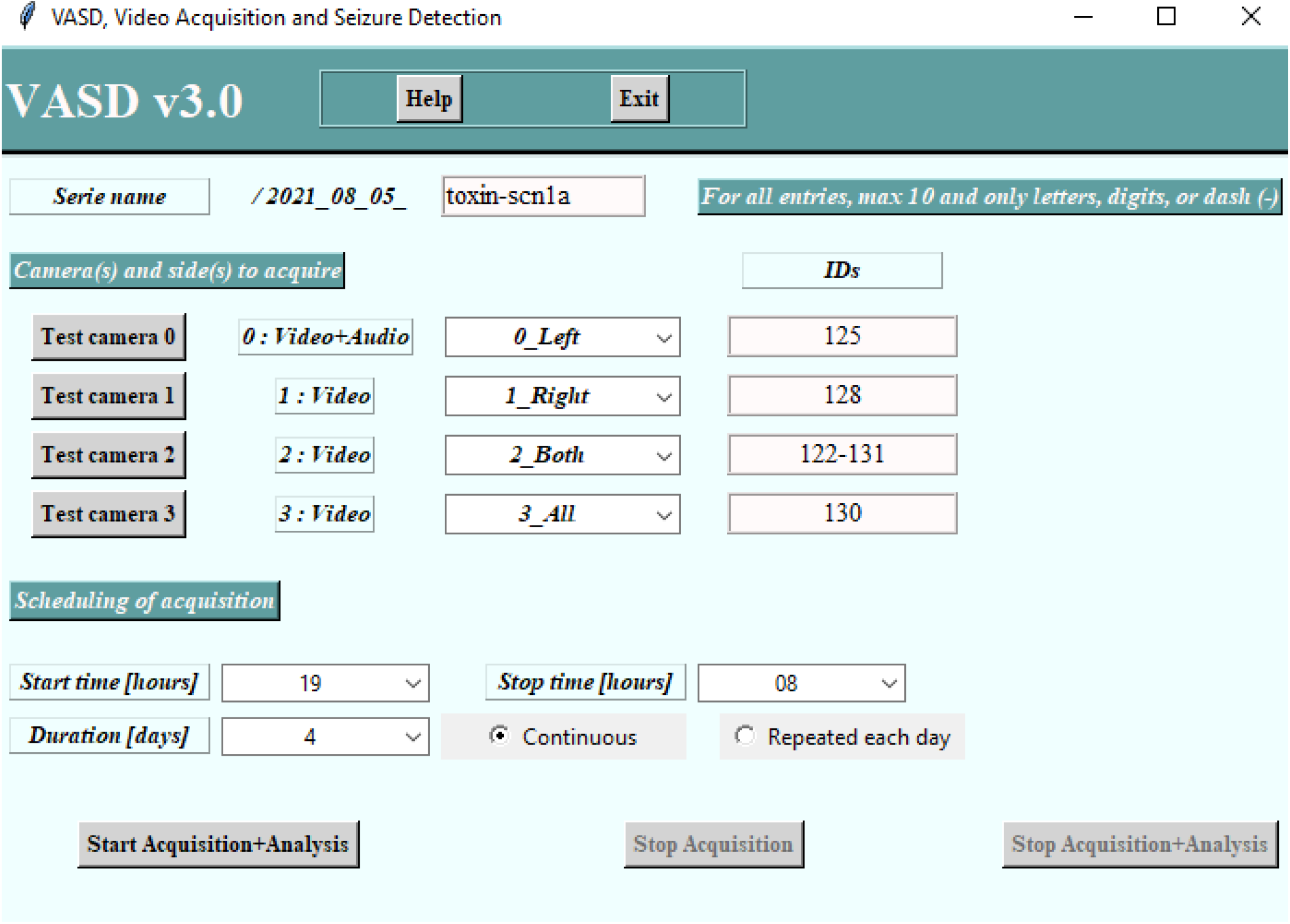
Example of VASD interface with the various acquisition options. User can visualize and select which camera and camera sides (left and right half-fields, or full field) to acquire and can name the serie and each camera, in this example we have online 4 cameras including one with a microphone (camera 0). The acquisition start and stop can be scheduled at any full hour and for up to 99 days. In the “Repeated each day” mode acquisition starts and stops every day at chosen hours (example: night acquisition between 20:00 and 8:00), otherwise acquisition is continuous. Acquired 1-hour long videos are stored as series, with one folder for each camera containing all corresponding video files.

#### 3.1.2 Automated motion detection and semi-automated seizure detection

SASDI (Semi-Automated Seizure Detection on Images) can run on window or linux machines (not tested on iOS). It enables to select lists of video files (any format accepted by opencv module) from any source (previously described VASD or other video acquisition systems) and with any length (tested up to 72 h long videos, but depending on file format 1h long videos are more efficiently analyzed). SASDI allows the selection of a full directory or a set of video files. It also proposes a very powerful selection mode: the serie. A serie is a folder containing other subfolders with video files (no theoretical limits in number; this structure is the one used by VASD for its output videos). SASDI enables the analysis in one go of all files within each subfolder with full field or user defined ROIs. It performs motion analysis with exactly the same algorithm as VASD. VASD and SASDI are optimized to analyze hundreds of files (tested up to 900 files in one go). This is particularly useful for analysis of long lasting chronic recordings. Moreover, SASDI add the possibility to draw and analyze separately up to 8 ROIs for each video within the same folder, which is a useful feature if the camera field contains more than one animal (e.g. several cages). VASD can analyze up to 2 ROIs automatically defined for each camera (left and right halves of the camera field), and many cameras can be recorded at the same time depending on the network bandwidth and computer performances (tested up to 5 cameras). To perform motion analysis, SASDI automatically used 80% of available threads but this value can be easily changed, in order to use all CPU threads, for maximizing the speed of analysis, or to use less threads, for being able to work at the same time with other software programs or not to overload a shared server. The parallel analysis of a multiple videos is very rapid (using one thread for each video in independent processes). For example, the analysis of 522 videos, recorded at 22 frames per seconds, on a server (Dell, CPU Intel Xeon Gold 5220) using only 10 threads and analyzing 2 ROIs of 640×720 pixels per video, was performed at 1090 Hz. The SASDI user interfaces for files selection and ROIs drawing are shown in Figure 4.

**Fig. 4.**
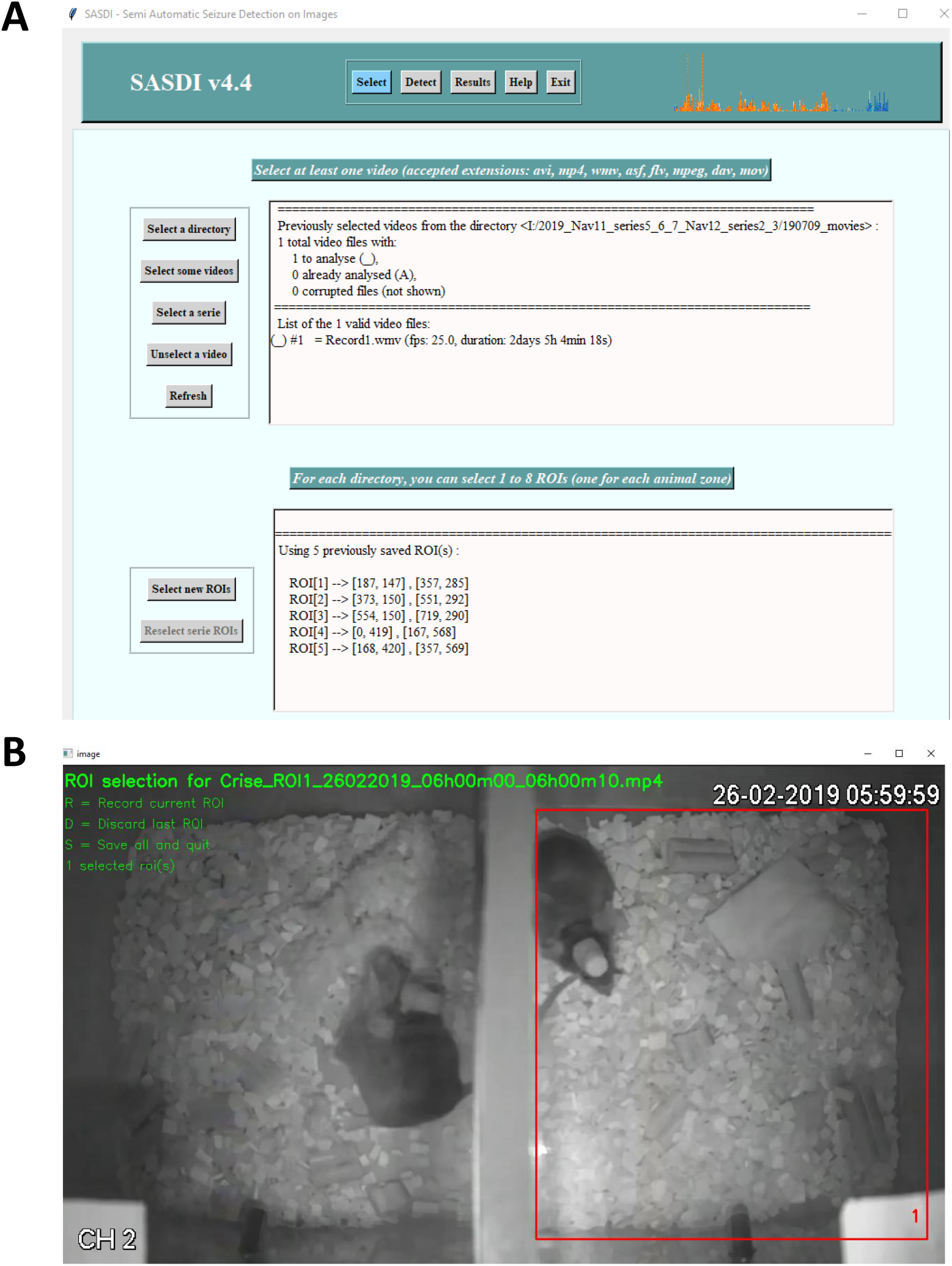
Example of SASDI interfaces with video files and animal selection. Screenshot of the “Select” page of SASDI interface (A). User can select a full directory, a selection of videos, or a serie (list of folders containing video files from VASD or from other sources), and draw on the first video image the required ROIs (B).

SASDI is also the reader for motion analysis files created by either VASD or SASDI itself (motion index against time, in csv format) and their corresponding video files. It permits to visualize on one graph the motion for each ROI of each video. It also enables to compute a spectrogram with consecutive Fourier transforms, displayed as the full bandwidth spectrogram every seconds (see above Algorithms paragraph). This analysis helps in the identification of convulsive seizures, because the frequency power quantified with the spectral analysis increases a lot during convulsive seizures. Note that this optional analysis is performed online, thus a fairly performant computer is needed. SASDI also enables to easily display the video at chosen time points to check that the detected motion peak is indeed a seizure. SASDI can also extract the audio stream from the video file, if present, and display it together with the quantified motion values. We added this option because certain animal models can emit a vocalization at the seizure onset, which can be used for seizure identification. An example of the results page of SASDI user interface and corresponding ECoG recording are shown in Figure 1.

SASDI is thus a tool that enables to scan rapidly hundreds of hours of videos recordings, especially useful for identifying convulsive seizures in rodents (grade 6 in most seizure severity scales).

### 3.2 Validation of motion analysis

#### 3.2.1 Usage of two murine epilepsy models

In order to validate the accuracy of the used algorithm we performed recordings on two murine models of genetic epilepsy that show spontaneous convulsive seizures. *SCN1A* (Na_V_1.1 sodium channel) mutations cause the severe developmental and epileptic encephalopathy Dravet syndrome (DS) and the in general milder Genetic Epilepsy with Febrile Seizure plus (GEFS+), as well as Familial hemiplegic migraine type 3 (FHM3) (Dravet 2011, Zhang, Burgess et al. 2017, Mantegazza and Cestele 2018, Mantegazza and Broccoli 2019, Mantegazza, Cestele et al. 2021). We used heterozygous *Scn1a* knock-out (*Scn1a*^+/-^) mice, which are an established animal model of DS, carrying a mutation that truncates Na_V_1.1 and showing hyperthermia-induced and spontaneous seizures, as well as behavioral-cognitive defects and increased mortality (Yu, Mantegazza et al. 2006, Liautard, Scalmani et al. 2013, Mantegazza and Broccoli 2019, Mantegazza, Cestele et al. 2021). The missense mutation R1648H causes mild GEFS+ in one family (Escayg, MacDonald et al. 2000) and DS in another family (Depienne, Trouillard et al. 2010). We used R1648H heterozygous knock-in (Scn1a^RH/+^) mice (Martin, Dutt et al. 2010, Hedrich, Liautard et al. 2014, Mantegazza and Broccoli 2019, Salgueiro-Pereira, Duprat et al. 2019, Mantegazza, Cestele et al. 2021), which in the mix C57Bl/6J-129/SV show a very mild phenotype that can be transformed into a severe DS-like phenotype after the induction of short repeated seizures (Yu, Mantegazza et al. 2006, Salgueiro-Pereira, Duprat et al. 2019).

#### 3.2.2 Behavioral characterization of seizures

The severity of seizures was quantified using the following scale (modified from Salgueiro-Pereira, Duprat et al. 2019): grade 0 - No behavioral response; grade 1 -Staring, 2 – Head nodding; grade 3 – Unilateral forelimb clonus; grade 4 - Fully developed bilateral forelimb clonus; grade 5 - Forelimb clonus with a tonic component and twist of the body; grade 6 - Convulsive seizures (clonic or tonic-clonic seizures with wild jumping).

#### 3.2.3 Seizure detection, comparison with human visual inspection

We performed an experiment with the knock-out Scn1a^+/-^ model recording only videos and with two animals per cage. An experienced experimenter visualized 351 hours of videos with 4 groups of 2 mice (4 controls and 4 Dravet syndrome model mice), and found 81 convulsive seizures (grade 6) and 49 milder seizures (scale 2 to 5). Another experienced user analyzed the same videos with SASDI and found 72 convulsive seizures out of the 81, representing an 88.8 % accuracy. Of note, the presence of two mice in the same cage is increasing the overall detected motion and masked some seizures. We also noticed that in some cases, the mouse experiencing a seizure was at the back of the cage while its littermate was in the front, partially hiding the former. This shows that recording from a lateral view and with two animals in the same cage is not optimal for image analysis, although even in these sub-optimal conditions SASDI had a good accuracy.

#### 3.2.4 Seizure detection: comparison with electrocorticogram (ECoG) recordings

The gold standard for seizure identification is the analysis of the recording of brain electrical activity. We acquired synchronized video and electrocorticogram recordings from R1648H *knock-in* mouse model with a total of 10 mice and 748 hours of recordings (daylight only). To improve image analysis animals were isolated, with one animal per half-cage (divided using a stainless steel separator with 1 cm holes to maintain physical contact and social interactions). ECoG recordings were visually analyzed by a first experimenter looking for the large abnormal hyperactivity characteristics of generalized convulsive seizures (see Figure 1B), each abnormal electrical activity was then checked on the corresponding video to confirm a behavioral seizure. With ECoG analysis we found a total of 71 seizures, of which 59 were convulsive (grade 6) and 12 were of minor severity (scale 2 to 5). A second experimenter analyzed only the video files using SASDI. Out of the 59 convulsive seizures, 55 were identified with SASDI, representing an accuracy of 93.2%. Examples of motion analysis with detected seizures are presented in Figure 1A. Note that it is very difficult to detect non convulsive seizures (in particular without wild jumping) using image analysis, because the animal motion during clonic or relatively mild tonic-clonic seizures is in the same range as common activities (walking, grooming, eating, …).

It has to be pointed out that we used standard videos normally used to evaluate behavioral features of seizures identified analyzing a synchronized ECoG signal recorded in parallel with a wired electrophysiological system; they are not optimal for image analysis for several reasons:

1. The recording system allow the acquisition of only one video stream together with ECoG acquisition. To optimize the animal usage, we usually record 8 mice in parallel, each in half a cage. Each cage is shot with one camera, and the 4-camera streams are concatenated into a single video stream of 720 x 576 pixels (total of 414720 pixels). Consequently, the analyzed ROI for each animal is about 180 x 140 pixels (25200 pixels), reducing the definition of the image and the precision of the analysis.
2. The videos were taken from a lateral view, when a mouse is moving from the front to the rear of the cage or vice versa its apparent size changes and the recorded motion is amplified.
3. The cable of the electrophysiological wired recording system induces unwanted motion artefacts, thus the chosen ROIs were smaller to avoid analyzing the space above the animals,

Thus, the comparison with the gold standard (analysis performed on the ECoG recording) shows a good accuracy for SASDI, although this value is underestimated because of the aforementioned problems and artefacts. However, the results obtained with the analyses of these sub-optimal videos further shows the robustness of SASDI.

#### 3.2.5 Activity, comparison with electromyogram (EMG) recordings

The gold standard for recording activity and especially wake/sleep phases is electromyogram (EMG), but this intrusive technique needs the implantation of electrodes in the neck muscle and a wire or wireless connection to the animal. The motion analysis provided by SASDI can provide a good assessment of animal activity. Figure 5 shows an example of EMG recording and the corresponding video motion analysis, the non-active phases are clear on both recordings and can also be used.

**Fig. 5.**
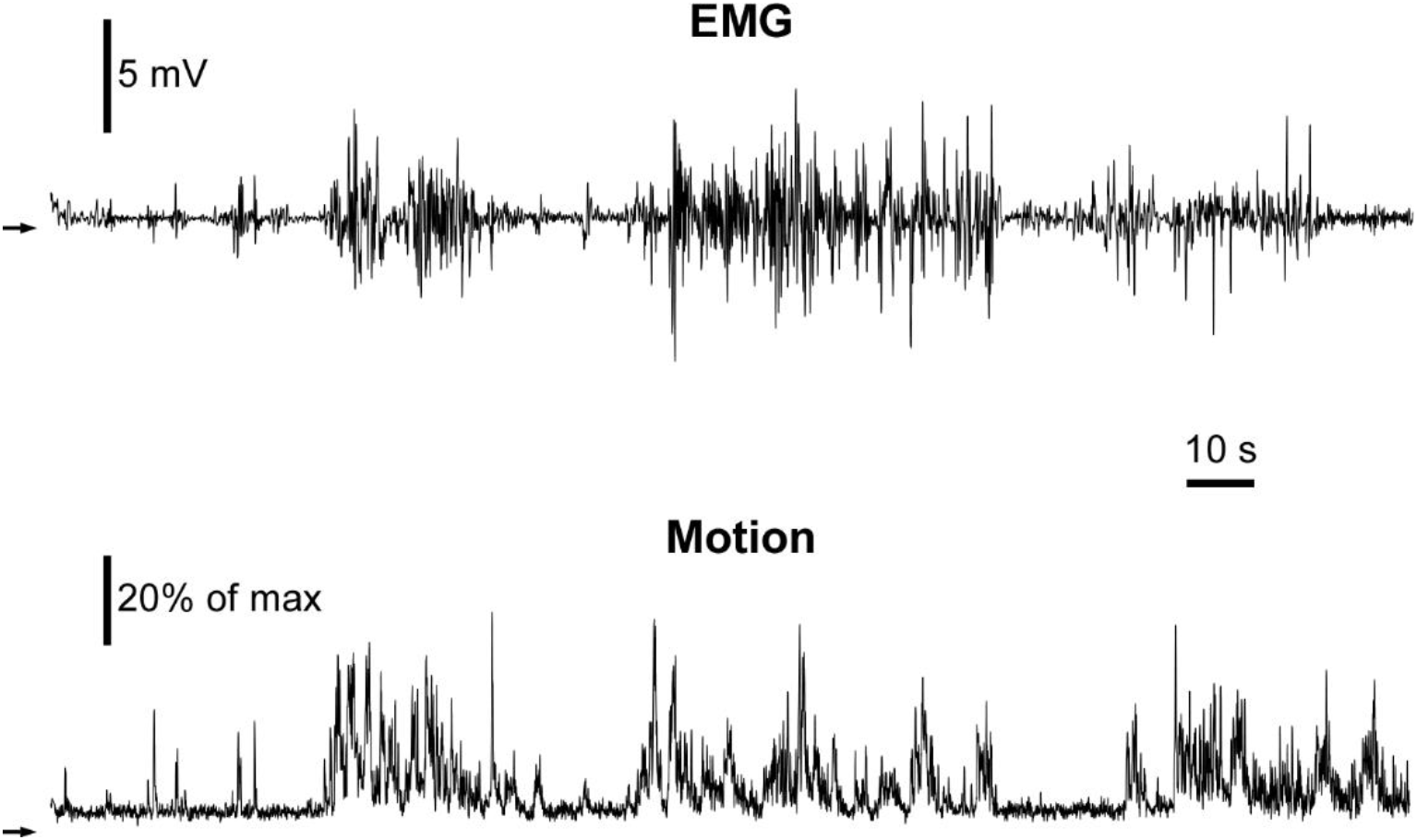
Animal activity from EMG and from motion analysis. Comparison of animal muscular activity measured with an electromyogram (EMG) recording showing recorded potential (mV) as a function of time (s) (top graph) and the corresponding motion analysis (in % of maximal motion) from SASDI video analysis (lower graph) the black arrows indicate 0 for each graph. Both graphs are clearly indicating the same overall mouse activity.

#### 3.2.6 Known bugs using VASD or SASDI

The video files container (mp4, asf, …) normally include headers with different types of information on the video (frame rate, duration, etc.). In some case, these informations are not properly recorded or the frame rate is variable, in that case SASDI will prompt the user to manually enter the frame rate. SASDI needs a constant frame rate to be able to properly calculate the time base; if it not the case, you can convert the file before analysis (using for example ffmpeg) or give an average frame rate (less accurate). Videos recorded with VASD are in the proper format.

#### 3.2.7 Installing VASD and SASDI

The two packages are available on GitHub (https://github.com/fabriceduprat/SASDI and https://github.com/fabriceduprat/VASD). All installations steps are detailed in each README files and a detailed User guide is available using Help buttons within each interface.

### 4 Discussion

Various commercial and open source software programs enabling detailed behavior analysis from video only are available. Nonetheless, none of them are designed to detect convulsive seizures on a large number of files.

Quantification of spontaneous seizures is essential for the characterization of rodent models of epilepsy, and the evaluation of therapeutic approaches on spontaneous seizures can greatly increase the translational value of the study (Loscher and Schmidt 2011). Convulsive seizures are observed in numerous epilepsy types (Scheffer, Berkovic et al. 2017) and they can lead to sudden unexpected death in epilepsy (SUDEP) (Massey, Sowers et al. 2014), which is a major concern for epileptic patients and caregivers. They are also observed in numerous rodent models, both of genetic and acquired epilepsy (Velíšková 2017).

VASD and SASDI is a free package that only needs cost-effective webcams and a computer to run. It allows the acquisition and analysis of hundreds of hours of video recording, providing high efficient identification of convulsive seizures and an overall index of the animal activity. This noninvasive approach allows the study of animals without any surgery and electrode implantation, and also allows to perform chronic studies of young animals, which are particularly difficult with implanted electrodes. In fact, we successfully used it for litters of pups aged 10 to 30 days (with their mother up to age 21 days); it is still a major gain of time compared to visual inspection of the full video recording, even if there are much more false-positive than with individual animals.

The main limitation of our image analysis system is that it cannot detect non convulsive seizures, because they are very close to normal activity. Another concern is the quality of video recording that can hamper the results.

To enhance video quality, we recommend to use the following advices before starting acquisition:

- Try to draw ROIs of equivalent surface to obtain a similar motion value for each animal.
- To avoid modifications of the observed size of mice, which in videos changes according to the camera-mouse distance, videos taken from above are better than from aside.
- To enhance contrast, use a dark background with light-colored animals, or vice-versa.
- To avoid misplaced ROIs, always keep the analyzed cage in the same place (add marks to reposition precisely the cage).
- For the same reason, be careful not to swap position of any elements, when changing cages (e.g. water dispenser, food dispenser, …).
- For better animal detection, use a homogenous illumination, not blinking, and with minimal shades, during both days and nights.
- For the same reason, check that there is no reflection on cages neither during night nor day illuminations, and that the camera does not produce artifacts at the chosen acquisition rate.
- To properly detect seizures, the best configuration is 1 animal per zone (use a separator with holes to allow social interactions). We showed that using pups with their mother the analysis is possible but is much more time-consuming.

In conclusion, the main advantages of using this new method for convulsive seizure detection in rodents, are 1) it is noninvasive, 2) the high speed of its analysis, with an average of 100 Hz per video file per CPU thread, 3) its ability to acquire and analyze a very large number of files with a good accuracy (above 88%), 4) its easy-to-use graphical interface, and 5) it is very cost effective.

## 5 CRediT author statement

Daniel Diaz-Arce: Software.

Anis Ghouma: Software. Paolo Scalmani: Investigation, Validation.

Massimo Mantegazza: Methodology, Resources, Writing-Review, Funding acquisition.

Fabrice Duprat: Conceptualization, Methodology, Software, Investigation, Validation, Writing-Review & Editing, Funding acquisition.

## 6 Competing financial interests

The authors declare no competing financial interests.

## 7 Acknowledgements

We gratefully thank Damien Barbier for the linux server settings. This research was supported in part by University Cote d’Azur “Académie d’Excellence 4” (https://univ-cotedazur.fr/structures-de-recherche/academies-dexcellence/academie-4/projets/projets-finances, Epi-Analyse), University Cote d’Azur UCA-JEDI (https://univ-cotedazur.fr/ucajedi-lidex-duniversite-cote-dazur, ANR-15-IDEX-01) and by the Laboratory of Excellence “Ion Channel Science and Therapeutics” - LabEx ICST (https://www.labex-icst.fr/en, ANR-11-LABX-0015-01).

## Notes

### Competing Interest Statement

The authors have declared no competing interest.

